# Plant-host shift, spatial persistence, and the viability of an invasive insect population

**DOI:** 10.1101/2021.09.20.461112

**Authors:** Isabelle Bueno Silva, Blake McGrane-Corrigan, Oliver Mason, Rafael de Andrade Moral, Wesley Augusto Conde Godoy

## Abstract

Assessing the effects of a plant-host shift is important for monitoring insect populations over long time periods and for interventions in a conservation or pest management framework. In a heterogeneous environment, individuals may disperse between sources and sinks in order to persist. Here we propose a single-species two-patch model that aims to capture the generational movement of an insect that exhibits density-dependent dispersal, to see how shifting between hosts could alter its viability and asymptotic dynamics. We then analyse the stability and persistence properties of the model and further validate it using parameter estimates derived from laboratory experiments. In order to evaluate the potential of this model, we applied it to *Drosophila suzukii* (Diptera: Drosophilidae), which has become a harmful pest in several countries around the world. Although many studies have investigated the preference and attractiveness of potential hosts on this invasive drosophilid, no studies thus far have investigated whether a shift of fruit host could affect such a species’ ecological viability or spatiotemporal persistence. The model results show that a shift in host choice can significantly affect the growth potential and fecundity of a species such as *D. suzukii*, which ultimately could aid such invasive populations in their ability to persist within a changing environment.

## 1. Introduction

The adaptation of non-native species to new environments is essential for biological invasion processes, which involve colonization, establishment, and local or regional dispersal events (Hengeveld 1989). Many invader populations have polyphagous habits, usually exploiting different host plants, which can result in these species becoming pervasive agricultural pests (Kennedy and Storer, 2000). Studies that focus on plant host shifts are important to elucidate patterns of movement in insects, so as to understand their dispersal dynamics and to provide vital information for integrated pest management (IPM) programs (Jian, 2019).

Mathematical models are simple tools commonly used to understand ecological patterns of the movement and population dynamics of invader species (Shigesada and Kawasaki, 1997.). In recent years, additional structure, be it spatial or demographic, has been incorporated into mathematical and computational models, as a first attempt to explain how insects move at different spatiotemporal scales and what various life stages contribute to their growth (Coutinho et al. 2012; Malaquias et al. 2017). Models that include spatial structure can offer a new perspective when investigating the movement of insects in patchy environments (Okubo and Levin 2001; Moretti et al. 2013).

Models of population growth combined with laboratory experiments can allow the dynamics of such organisms to be simulated over many successive generations based on empirical parameter estimates, longer than what is feasible in vivo (Heaps et al. 2016; Dey and Joshi 2018). Structural models that have been proposed in the past have tried to model the dynamics of insect species by incorporating, for example, probability of occurrence (Ørsted & Ørsted 2018) and the influence of abiotic factors (Langille et al. 2016; Reyes & Lira-Noriega 2020; Winkler et al. 2020), however to our knowledge no model has tried to assess the effects of a host-shift on a species’ persistence.

In the present study, we investigated how host shift may affect the local establishment, equilibrium dynamics, and persistence of such a species by combining a laboratory experiment using the newly invasive pest *D. suzukii*, and a mathematical model of dispersal. This multidisciplinary approach is useful to understand how insects may respond to variations in host availability, which for example may promote or hinder the ability of such animals to persist in new multi-crop landscapes. Our main objectives were to (1) assess the possible effects of host shift on the equilibrium dynamics and persistence of an insect population using a two-patch dispersal model, and to (2) parameterise the model via experimentally validated estimates of survival and fecundity for *D. suzukii*.

## 2. Methods

### 2.1. The model and local patch dynamics

To assess the possible effects that shifting between hosts can have on a population’s abundance we first propose a two-patch mathematical model that incorporates density-dependent dispersal and resource heterogeneity. Dispersal is, in general, an asymmetric process and may have significant consequences for population persistence (Rinnan 2018), especially when assessing the impacts of climate change and monitoring species range shifts (Zhou and Kot 2011). When introducing explicit spatiality into population models, a phenomenon known as Turing instability may arise, whereby the extinction equilibrium in the absence of dispersal is asymptotically stable, but by allowing dispersal it becomes unstable (Neubert et al. 2002). Thus, including spatial structure in population models may reveal how such instabilities arise and indicate the mechanisms that drive dispersal.

Consider a polyphagous, single-species population that inhabits two nonidentical patches, and denote the patch population densities, in generation 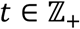, by *x*_1_(*t*) and *x*_2_(*t*), respectively. Following reproduction on a patch we assume both subpopulations disperse between patches, which in turn depends continuously on their respective densities (Figure 1). We also assume that the overall population is spatially closed, i.e., individuals reproduce and disperse only on the two specified patches. This may not be ecologically realistic, but our aim is to understand the potential effects of dispersal, or host shift, between two differing patches on the overall population dynamics. A general form for such a model is thus given by the system of difference equations

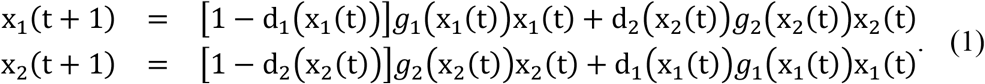

**Figure 1.**
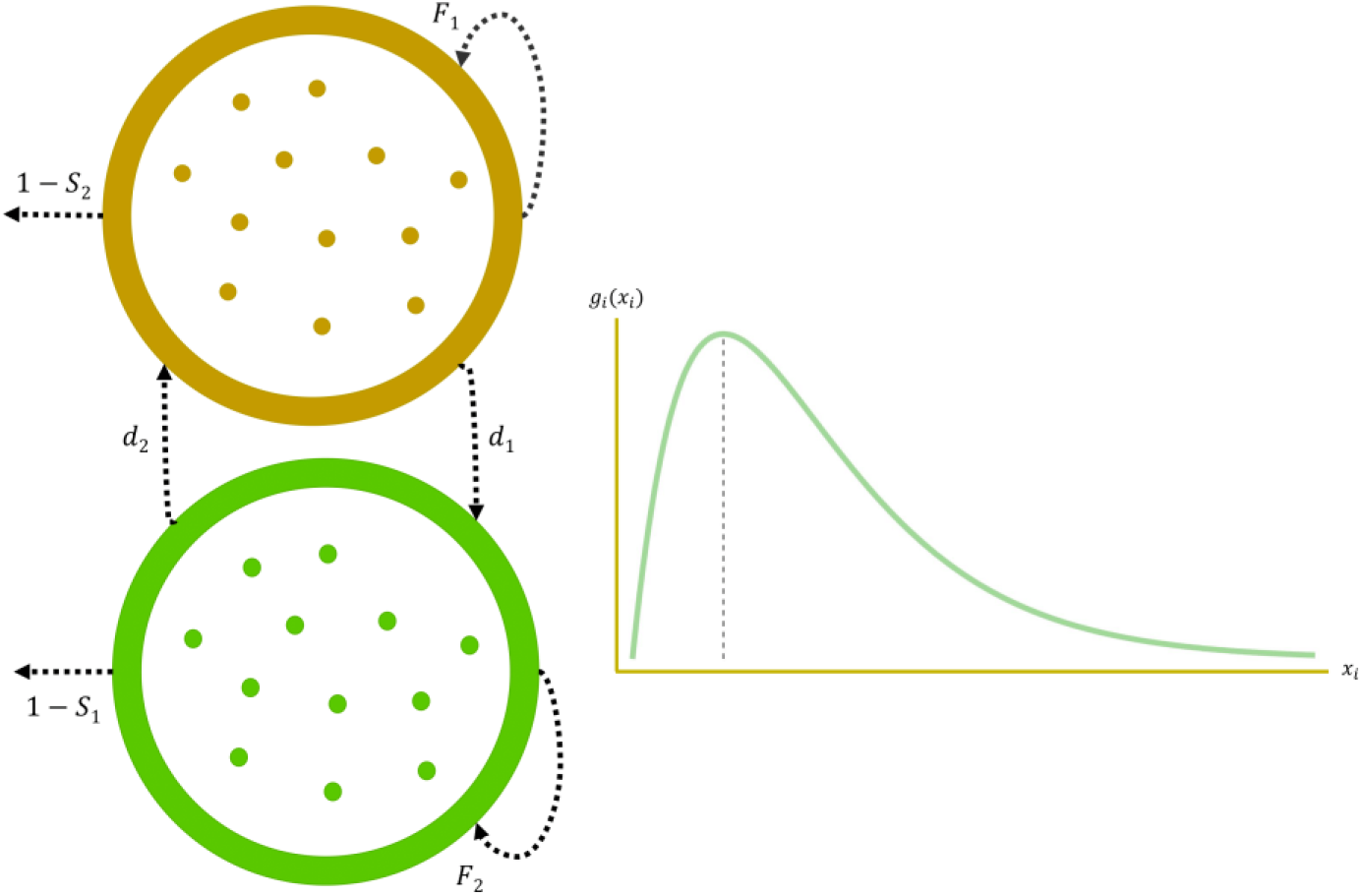
Population dynamics within and between hosts (left). *F_i_* and *S_i_* are respectively the fertility and survival proportion on patch i. Density-dependent dispersal from patch *i* is described by the function *d_i_*. Local dynamics (right) for patch *i* is described by the unimodal function *g_i_*.

This formulation considers two continuously differentiable functions, 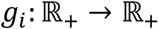 and 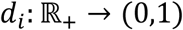, that respectively describe local growth on patch *i* and density-dependent dispersal from patch *i*. We assume dispersal is asymmetric, *d_i_* ≠ *d_j_* for *i* ≠ *j*, and bidirectional, *d_i_* ∉ {0,1}. Later, we will briefly discuss the case when dispersal is not permitted in either direction, i.e., each patch is isolated. We can rewrite (1) in nonlinear matrix model form

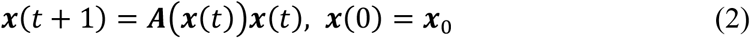

where ***x*** = (*x*_1_ *x*_2_) is a state vector in 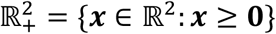, the cone of 2-dimensional real vectors with nonnegative components, and 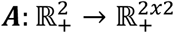 is a continuous function that maps nonnegative vectors into nonnegative square matrices, commonly written in matrix form (Cushing 1998). Our nonlinear population projection matrix is thus given by

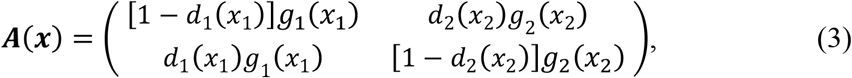

where *d_i_*(*x_i_*) ∈ (0,1) for all *x_i_* ≥ 0, *i* = 1,2. Typically, matrix projection models are often considered when populations show clear stage or age structure (Smith and Thieme, 2011), for example when using Leslie matrix models (Leslie 1945), but we will use them to partition our population into two patches, each patch with its own vital rates.

Before analyzing model (2), with ***A***(*x*) given by (3), we will recall some key mathematical definitions needed for the results described later. For more detail on these and related results, the reader should consult the Supplementary Material.

An equilibrium ***x***^⋆^ of (2) is a solution of ***x*** = ***A***(***x***)***x***. The equilibrium ***x***^⋆^ is said to be stable if for any *ϵ* > 0 there exists a *δ* > 0 such that |***x***_0_ – ***x***^⋆^| < *δ* implies that |***x***(t) – ***x***^⋆^| < *ϵ* for all t, where ***x***_0_ denotes the initial condition ***x***(0). If, in addition, there is some *R* > 0, such that ***x***(*t*) → ***x***^⋆^ as *t* → ∞ for any solution with |***x***_0_| < *R*, the equilibrium is locally asymptotically stable. If this holds for any *R* > 0, it is said to be globally asymptotically stable. Clearly, any nonlinear matrix model has an equilibrium at the origin, aka the extinction equilibrium. If the extinction equilibrium is (locally or globally) asymptotically stable, then lim_*t*→∞_***x***(*t*) = **0**, i.e. the state vector asymptotically tends toward the origin. The mathematical definitions of persistence, which we will now recall, characterize the opposite situation, where populations are maintained at some positive level asymptotically.

Given a general nonlinear matrix model of the form (2) with state space 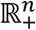 we say that this nonlinear system is uniformly weakly persistent with respect to a persistence function 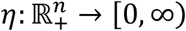 if there exists some *ϵ* > 0, such that for each solution ***x***(*t*) = (***x***_1_(*t*) ***x***_2_(*t*) ⋯ *x_n_*(*t*)), with initial condition 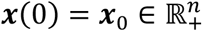 and *η*(***x***_0_) > 0,

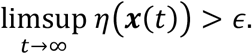

The persistence function is typically some quantitative measure of population abundance, such as the total or site-specific population size for example. Intuitively, weak uniform persistence means that for any time *T* > 0, there is some *t* > *T* at which the persistence measure exceeds *ϵ*. A stronger concept of persistence is that of uniform (strong) persistence, which is identical to the above definition apart from replacing the *limsup* with the *liminf*. Strong uniform persistence means that there is some *T* > 0 such that the persistence function exceeds *ϵ* for all later times *t* > *T*.

We will assume that the local growth function takes the form

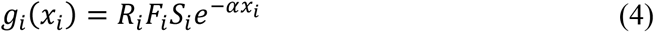

where *F_i_* > 0, *S_i_* ∈ (0,1), *R_i_* ∈ (0,1) and *α* > 0, for *i* = 1,2 (see Figure 1). We also refer to (4) as the recruitment or fitness function (Chesson 2012; Eager at al. 2012). Parameters *F_i_*, *S_i_* and *R_i_* can be interpreted as respectively being the fecundity, survival proportion and sex ratio on patch *i*. Density-dependence is assumed to affect each local population in the same way, with *α* being the influence of intra-specific competition for resources. Note that the difference in each patch is determined by the effects of nutrition and fecundity, as expressed through *S_i_* and *F_i_* being different for each patch, i.e., resource heterogeneity. These factors ultimately determine if each generation successfully reproduces and survives over each time step.

Recruitment function (4) was inspired by Prout and McChesney (1985), who used a difference equation to describe intraspecific competition among an immature *D. melanogaster* population and ultimately how this affected adult fertility and survival. In our model we use constant demographic parameters and describe the influence of patch specific density-dependence through *e*^−*αx_i_*^. When the two patches are isolated, the dynamics on patch *i* is given by

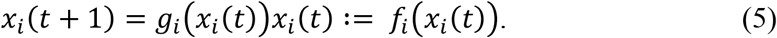

This is similar in form to a Ricker map (Ricker 1954). It is unimodal, meaning it attains a unique highest value or global maximum (see Figure 1). This occurs at *x_i_* = *α*^−1^. The maximum possible population size is then

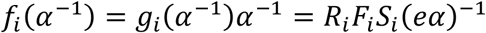

and so, the population is prohibited from reaching arbitrarily large values. It can also be seen from (4) and (5) that if *R_i_F_i_S_i_* < 1, for *i* = 1,2, then *x_i_*(*t*) → 0 as *t* → ∞. A unique positive equilibrium for patch *i*, 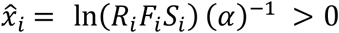 if, and only if, *R_i_F_i_S_i_* > 1. *f_i_* is continuously differentiable at 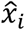, meaning the local stability properties of this equilibrium can be determined by the linear approximation given by

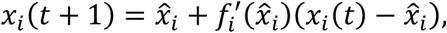

provided the difference 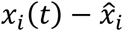 is sufficiently small for all *t* ≥ 0 (Elaydi, 2005). Local asymptotic stability is guaranteed once 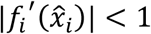. Thus 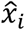 is locally asymptotically stable precisely when

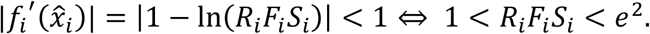

### 2.2. The assay

We reared a population of *D. suzukii* from colonies maintained in the Entomology Laboratory of the Federal University of Pelotas, Rio Grande do Sul State, Brazil. The flies used for the assays were kept in cages (30 × 21 × 17 cm) in climate chambers (Fitotron®) at 25 ± 1°C, relative humidity 70 ± 5% and 14–10 h photoperiod. *D. suzukii* was chosen as a model insect in the present work because it is a continental invasive species of high relevance for fruit growing worldwide and especially in Brazil. The flies received fresh raspberries cultivated without insecticides, obtained from farms in southeastern Brazil, to obtain the first generation. Fruits with eggs were removed and incubated under the same conditions until emergence. Raspberries were chosen as hosts as they are subject to high fruit-fly infestations in Brazil (Klesener et al. 2018). Flies that emerged in raspberry (generation F1) were placed on either raspberry or strawberry, with five replicates, in a completely randomised design (Figure 2). Each replicate consisted of one test tube (7.8 cm height × 3.8 cm diameter) with one strawberry or two raspberries and ten couples of *D. suzukii*. After the adults of the next (F2) generation emerged, they were placed in another test tube containing raspberries or a strawberry. This step was repeated, in a completely randomised design, but now with three replicates, each consisting of a test tube with five females and three males of *D. suzukii* from the F2 generation (Figure 2). Then, we repeated the entire assay using the same protocols, with the only difference being the number of replicates for the F2 generation, which was five for the second assay (Figure 2). The fruits were replaced daily, and the number of eggs and daily mortality of adults were recorded over the F1 and F2 generations. We observed the total number of eggs laid in each replicate, egg-adult viability, total oviposition period (in days), and survival time of the adults (in days).

**Figure 2.**
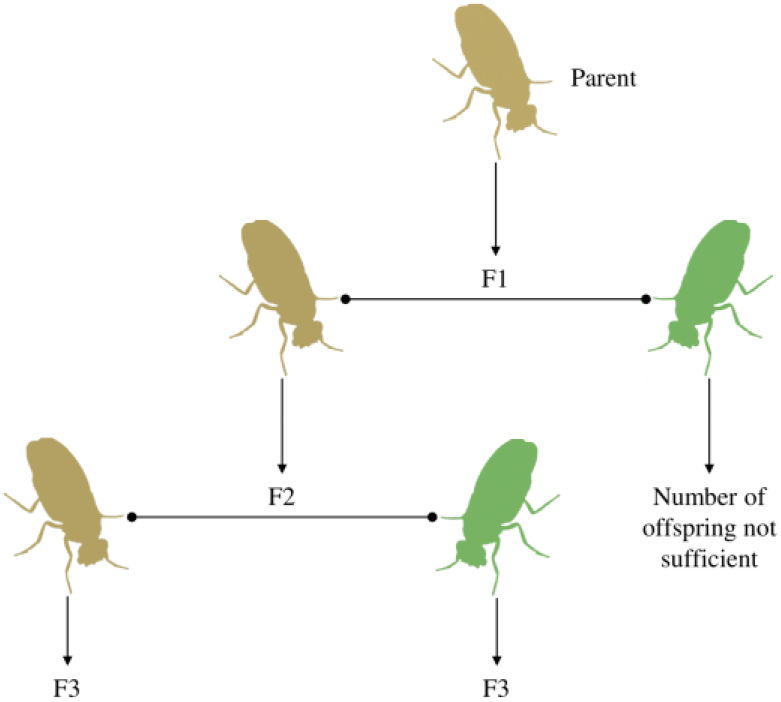
Host-shift assay showing the experimental design performed. For the raspberry filial generation reared on strawberry there were insufficient population numbers to produce a second strawberry filial generation.

**Figure 3.**
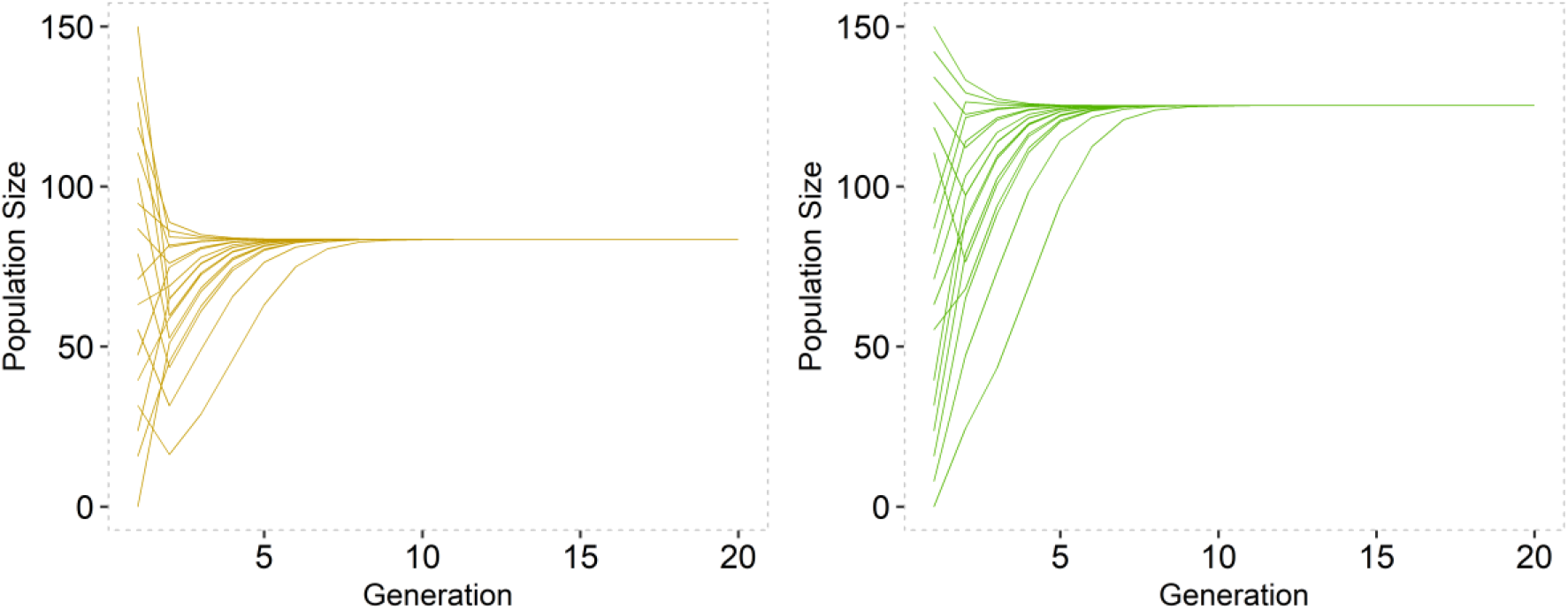
Patch trajectories for patch 1 (left) and patch 2 (right) with R_i_F_i_S_i_ given by empirical estimates obtained from experiments and constant/passive dispersal parameters d_1_ = 0.6 and d_2_ = 0.4. They approach stable positive equilibria for all initial conditions between 1 and 150.

### 2.3. Analysis of macronutrient concentrations

Three randomly selected adults from each generation (F1, F2 and F3, Figure 1) were placed individually in Eppendorf tubes containing 100% ethanol. Four macronutrients: protein, lipid, carbohydrate and glycogen were extracted from each fly, following Lorenz’s (2003) protocol, and prepared separately for quantification in a spectrophotometer (Epoch Microplate Spectrophotometer, BioTek).

Protein was quantified using 300 μL of Coomassie reagent (Coomassie Plus Protein, Pierce Biotechnology, Inc., Rockford, IL, USA), and 10 μL of each sample, in triplicate. Five standard curves were prepared, based on concentrations from 0 μg/mL to 2000 μg/mL, following the manufacturer’s protocol. The absorbance used was 595 nm. For lipid quantification, we used vanillin as the reagent in a sulfuric acid solution, and 10 μL of each sample, in triplicate. We used commercial vegetable oil to obtain the five standard curves (concentrations of 0 mg/dL to 2500 mg/dL), at an absorbance of 525 nm (Van Handel 1985). Carbohydrates and glycogen were quantified using anthrone reagent (Roe 1955). For the carbohydrate aliquots, we prepared a 50 μL solution of each sample individually, containing 100 μL of sulfuric acid and 200 μL of anthrone. The glycogen aliquots were prepared with a solution containing 25 μL of the sample, 50 μL of sulfuric acid and 100 μL of anthrone. The five standard curves were constructed, with glucose used as default (concentrations of 0 mg/dL to 100 mg/L) and an absorbance of 620 nm.

### 2.4. Statistical analyses

Quasi-Poisson models (Demétrio et al. 2014) were fitted to the data for total number of eggs, including the effects of assay (to reflect the experimental design), generation, host, and the interaction between generation and host in the linear predictor, and the natural logarithm of the number of females as an offset term. F tests were performed to assess the significance of the effects, and multiple comparisons were performed by obtaining the 95% confidence intervals for the linear predictors. Goodness-of-fit was assessed using half-normal plots with simulation envelopes (Moral et al. 2017).

To study the association between oviposition behaviour over time and survival times of the insects, a joint model for longitudinal outcomes (oviposition) and time-until-event outcomes (survival times) was fitted to the data (Rizopoulos 2010). For the longitudinal part of the model, we log-transformed the count data and included the effects of the assay (to reflect the experimental design), generation, host, time, all two-way interactions between generation, host, and time, and the three-way interaction amongst these factors in the linear predictor as fixed effects. We also included a random intercept per female, given that the number of eggs laid by the same female is correlated over time. For the survival part of the model, we used a Cox proportional-hazards model and included the effects of the assay, generation, and host, and the interaction between generation and host in the linear predictor. This modelling strategy allows for the estimation of an association parameter, which in this study measures the effect of fecundity on the mortality risk (Rizopoulos 2010). We performed likelihood-ratio (LR) tests to assess the significance of the fixed effects for both parts of the model and a Wald test to assess the significance of the association parameter.

The quantitative data for metabolised macronutrients were analysed by fitting linear models including the effect of treatment (F1 raspberry, F2 raspberry, F3 raspberry, and F3 strawberry) in the linear predictor. The significance of the treatment effect was assessed using F tests.

## 3. Stability and persistence results

We will now consider model (1) rewritten as a nonlinear matrix model (2), with ***A***(***x***) given by (3) and local patch dynamics given by (4). We assume that dispersal is bidirectional, asymmetric and a density-dependent process. In the rest of the paper, *ρ*(***B***) denotes the spectral radius of a square matrix ***B***. We refer the reader to the Supplementary Material for formal definitions and proofs of the results presented below.

As a first example, let us look at when we have passive/constant dispersal, i.e., *d_i_*(*x_i_*) = *p_i_* ∈ (0,1) for all *x_i_* ≥ 0. As *e*^−α*x_i_*^ ≤ 1 for *x_i_* ≥ 0, *i* = 1,2, it can be seen that ***A***(***x***) ≤ ***A***(**0**) for all ***x*** ≥ 0 in 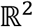. It follows that for any 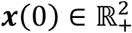,

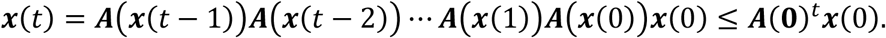

So, we have that if *ρ*(***A***(**0**)) < 1, then ***A***(**0**)^*t*^ → 0 as *t* → ∞. Thus, the extinction equilibrium is globally asymptotically stable. This gives a sufficient condition for ensuring patch extinction, as is shown in Smith and Thieme (2011) for general nonlinear matrix models.

In the context of pest management, we are primarily interested in how the presence/absence of dispersal influences persistence over time and what drives the population on either patch to extinction (eradication). We are also interested in knowing if on either patch, both assumed to differ in nutritional quality, whether the subpopulations persist even at low densities. This is a major concern in agricultural management, namely small numbers of individuals persisting on ephemeral low-quality habitat and then exhibiting rapid growth when favorable conditions arise (Reigada et al. 2018).

It is well known that for any induced matrix norm | · |, the spectral radius of ***B*** satisfies *ρ*(***B***) ≤ |***B***| (Horn and Johnson, 2012). As the *l*_1_ induced matrix norm is given by the maximum of the column sums of a matrix, it follows that ρ(**A**(**0**)) <1 if all column sums of **A**(**0**) are less than 1. This implies that a sufficient condition for ρ(**A**(**0**)) <1 is R_i_F_i_S_i_ < 1. This could be interpreted as each patch having either low survival proportion and fertility, high fertility but low survival probability or vice versa, which intuitively will lead to the eventual decline of the population in the long run.

For the remainder of this section, we consider what happens when *ρ*(***A***(**0**)) > 1, so local asymptotic stability of the extinction equilibrium is not guaranteed. The results presented below, will show that dispersal can lead to patch persistence when its absence could result in extinction on one of patches. This is akin to dispersal driven growth or Turing instability (Neubert at al. 2002; Guiver et al. 2017), whereby allowing dispersal prevents a previously declining and isolated population from going extinct, a somewhat similar concept to the so-called rescue effect in metapopulation theory (Hanski 2008).

We will first note that *G*(***x***) ≔ **A**(***x***)***x***, with **A**(***x***) given by (3) and local patch dynamics given by (4), is strongly positive. Here, the notation ***x*** ≫ **0** means that all components of the vector ***x*** are positive.

### Lemma 1.

Let ***A***(***x***) be given by (3) and local patch dynamics given by (4). Then for any 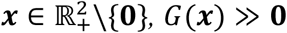.

Lemma 1 also implies that 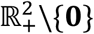 is forward invariant under *G*, i.e., trajectories that begin inside the positive orthant remain there for all time. In our next result, we note that when the spectral radius of ***A***(**0**) is greater than 1 there exists a positive patch-coexistence equilibrium of (2), satisfying *G*(***x***) = ***x***.

### Proposition 1.

Let ***A***(***x***) be given by (3) and local patch dynamics given by (4). Suppose that *ρ*(***A***(**0**)) > 1. Then there exists some ***x*** ≫ **0** with *G*(***x***) = ***x***.

In the following lemma, we show that for ***A***(***x***) be given by (3) and local patch dynamics given by (4), the system (2) is point dissipative. This simple fact is needed for the results on persistence that follow.

### Lemma 2.

Let ***A***(***x***) be given by (3) and local patch dynamics given by (4). Then there exists some *M* > 0 such that for any 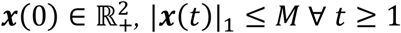.

The previous lemma is needed to establish the first of our main results.

### Theorem 1.

Let ***A***(***x***) be given by (3) and local patch dynamics be given by (4). Suppose that *ρ*(***A***(**0**)) > 1. Then, there exists some *ϵ* > 0 such that

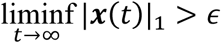

for any solution with 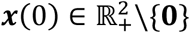.

This means that when *ρ*(***A***(**0**)) > 1 system (2), with nonlinear projection matrix given by (3) and local patch dynamics given by (4), is uniformly strongly persistent with respect to the total population size across the two patches. We will next show that a stronger result than Theorem 1 is possible, as *G*(***x***), with ***A***(***x***) given by (3) and local patch dynamics given by (4), is strongly positive.

### Theorem 2.

Let ***A***(***x***) be given by (3) and local patch dynamics given by (4). Let 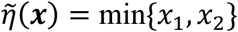 and suppose that *ρ*(***A***(**0**)) > 1. Then, there exists some *ϵ* > 0 such that

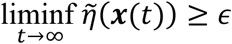

for any solution with 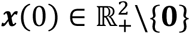.

As a second example, let us look at the scenario when *R*_1_*F*_1_*S*_1_ < 1 and *R*_2_*F*_2_*S*_2_ > 1, so on patch 1 the population tends to extinction and on patch 2 there is high survival and fertility. For *d_i_*(0) ∈ (0,1) we have that

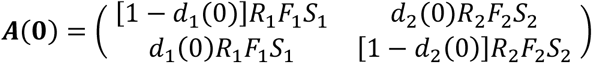

and so, for *R*_2_*F*_2_*S*_2_ sufficiently large, one can ensure that both uniform strong persistence and patch-persistence occur. For example, this can be easily seen if we let *R*_1_*F*_1_*S*_1_ = 0.9, *R*_2_*F*_2_*S*_2_ = 50, *d*_1_ = 0.5 and *d*_2_ = 0.7, meaning that *ρ*(***A***(**0**)) ≈ 16.01 > 1. By Theorem 2, this implies that, even though on one patch extinction may be inevitable, by allowing dispersal one cannot only rescue the declining patch from extinction, but also ensures that each individual patch population persists.

Locally a sink population may become extinct in the absence of immigration or source patches, but dispersal can allow the overall population the opportunity to persist. For fast-growing species with short life cycles and for those that exhibit irruptive dynamics, like for example insect species, populations can reach (and exceed) their carrying capacity very quickly, likely leading to high intraspecific competition and density dependent dispersal (Powell and Bentz 2014; Goodsman et al. 2017). Low quality habitat (sinks) that are distributed among higher quality habitat (sources) can limit the movement of organisms in space, and so identifying the mechanisms that limit growth can vary between habitat types (Heinrichs et al. 2016). Our simple modelling approach suggests that dispersal and demographic effects are potential drivers of population persistence. It is for this reason that modelling population dynamics within a spatial framework can provide new qualitative insights into the mechanisms that drive source-sink dynamics.

## 4. Experimental results

### 4.1. Total fecundity, oviposition period and proportion of viable eggs

The daily oviposition rates per female over time are shown in Figure 4. The interaction between generation and host significantly affected the total number of eggs (*F*_1,32_=11.05, p=0.0023). The number of eggs laid by females was higher in raspberry than in strawberry, for F1 and F2 generations; and the number of eggs laid in strawberry was higher in generation F2 than in F1 (Table 1). When setting adults of F1 generation in first host shift (Figure 2), available eggs were insufficient to have adults and to set up the second host-shift, as performed with raspberry. The interaction between host and generation did not significantly affect the total oviposition period (LR = 0.54, p = 0.4606). The global test for risk proportionality suggested that the model well fitted the data (*χ*^2^ test statistic = 3.16, d.f. = 4, p = 0.5316). Thus, analysing the marginal means indicated that the mean period of oviposition did not differ between generations (LR = 0.10, d.f. = 1, p = 0.7473), but did differ between hosts (LR = 31.77, d.f. = 1, p < 0.0001). Hence, females spent more time ovipositing in raspberry than in strawberry (Table S1 in Supplementary Material).

**Figure 4.**
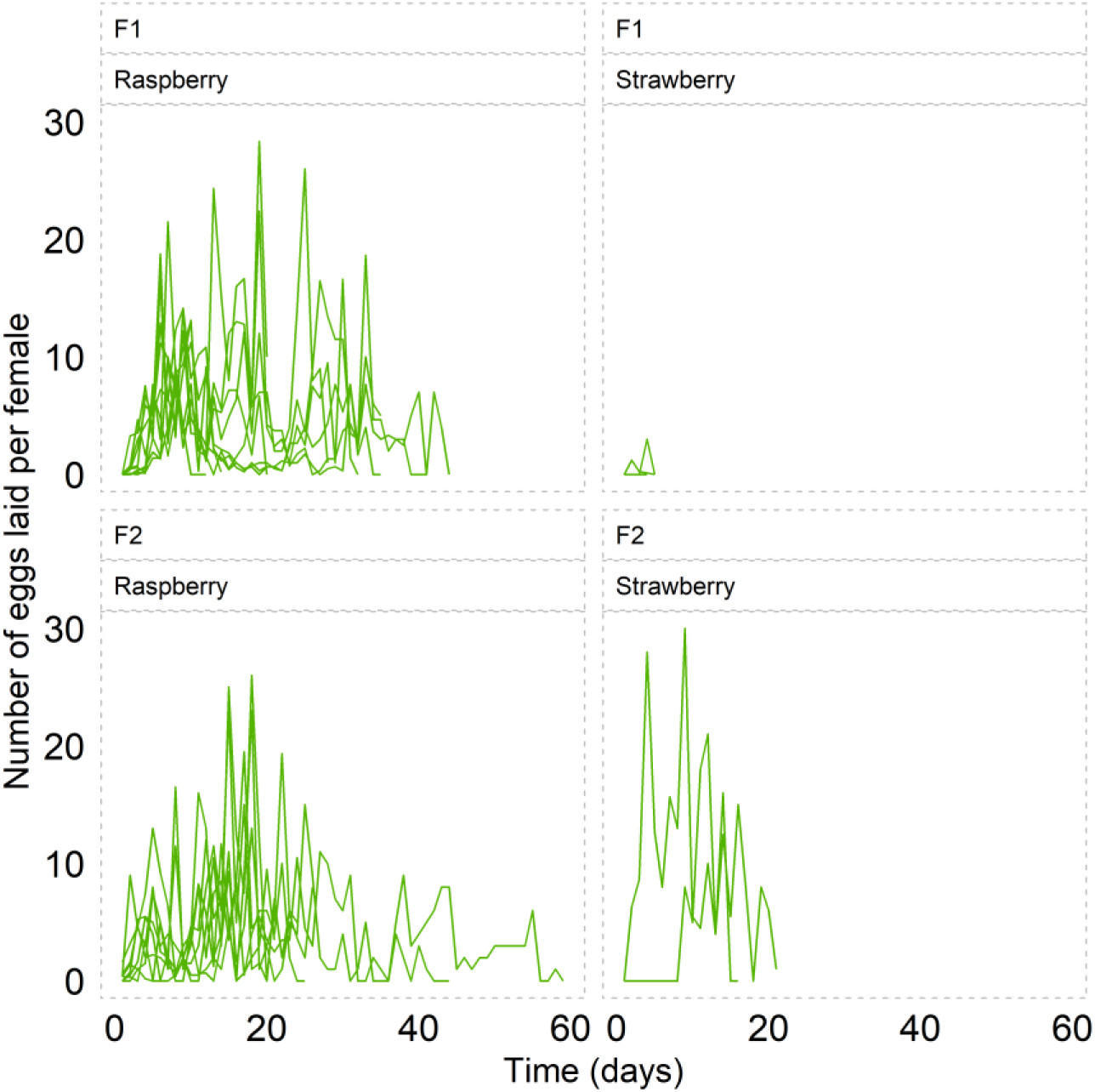
Number of eggs laid per female, in the F1 (A and B) and F2 (C and D) generations. Each line represents one replicate, which included ten and five females for generations F1 and F2, respectively.

**Figure 5.**
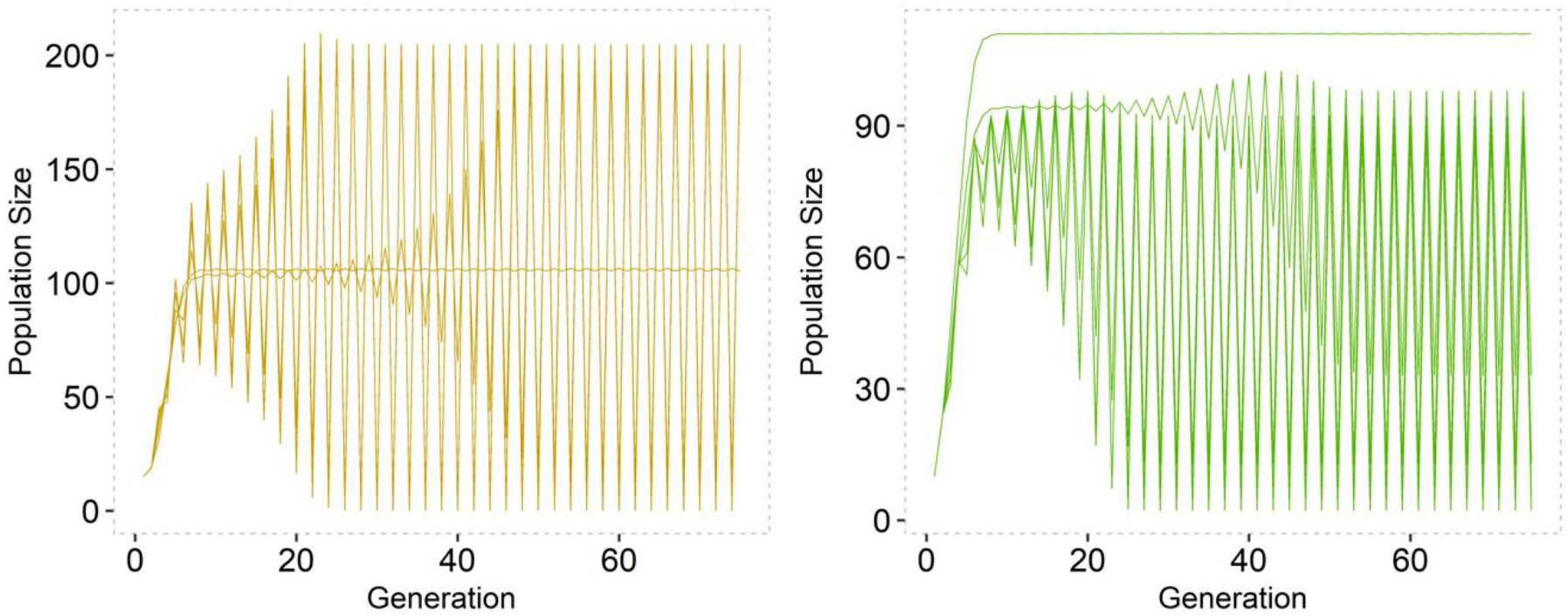
Boost-bust type dynamics for both patch 1 (left) and patch 2 (right), where *R_i_F_i_S_i_* are given by empirical estimates and *μ_i_* varies from 0.01 to 0.05. Initial conditions were (10,10). After initially increasing, trajectories either approach stable equilibria or oscillate indefinitely.

**Table 1:** Mean fecundity ± standard error (number of eggs laid per female), for each host and generation. Different upper-case letters denote significant differences in the rows; different lower-case letters denote significant differences in the columns.

| Generation | Host |  |
| --- | --- | --- |
|  | *Raspberry* | *Strawberry* |
| *F1* | 67.95 ± 7.70 aA | 0.20 ± 0.15 bB |
| *F2* | 50.40 ± 6.44 aA | 12.98 ± 11.03 aB |

The interaction between generation and host did not significantly affect the proportion of offspring that reached the adult stage (*F*_1,18_ = 0.07, p = 0.8004). However, there were significant main effects of host (*F*_1,19_ = 6.88, p = 0.0172) and generation (*F*_1,20_ = 25.77, p < 0.0001). Even though more eggs were laid in raspberry than in strawberry (see Table 2), the offspring were significantly more viable in strawberry (Table S2 in Supplementary Material).

**Table 2.**
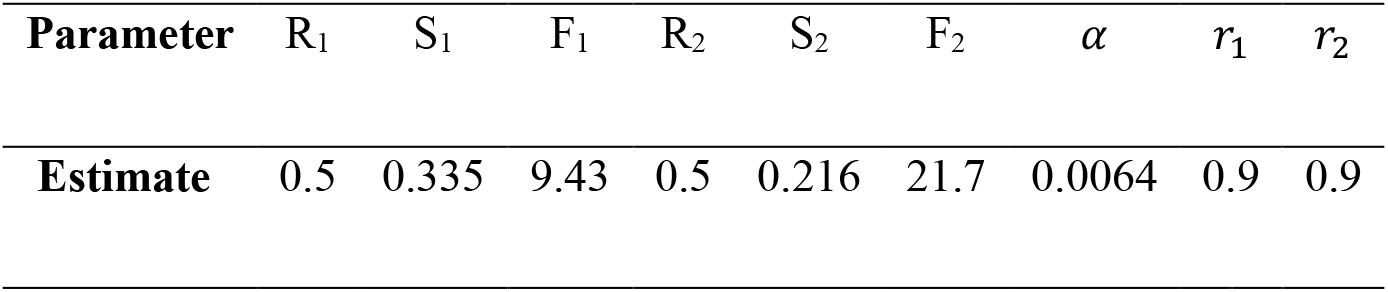
Demographic and dispersal parameters used in bifurcation plots and simulations. R_1_ and R_2_ were assumed to be equal. S_i_ and F_i_ were respectively the average number of eggs laid per individual and average number of individuals surviving on one host. An estimate for α was obtained from Prout and McChesney (1985). r_1_ and r_2_ were assumed to be equal.

### 4.2. Fecundity-survival time relationship and analysis of macronutrients

The results from the joint modelling of longitudinal oviposition and survival time indicated no significant effect of the three-way interaction between time, generation, and host (LR = 0.81, d.f. = 1, p = 0.3678), nor of the two-way interactions between generation and time (LR = 1.54, d.f. = 1, p = 0.2145) or host and time (LR = 2.4, d.f. = 1, p = 0.1210). However, the interaction between generation and host was significant (LR = 6.53, d.f. = 1, p = 0.0106), as was the main effect of time (Wald test p < 0.0001). Since the time estimate was negative (−0.0197 with a standard error of 0.0043), we concluded that the oviposition rate decreased over time. For the survival part of the model, the interaction between generation and host was not significant (LR = 0.17, d.f. = 1, p = 0.6777), nor was the main effect of generation (LR = 1.04, d.f. = 1, p = 0.3073). However, the main effect of host was significant (LR = 6.82, d.f. = 1, p = 0.0090), which suggests that insects reared on raspberry survived longer. We conclude that high fecundity is associated with a lower mortality risk, i.e., a longer survival time on the fruit, given the negative estimate of the association parameter (−0.5608 with a standard error of 0.1476, Wald test p < 0.0001). The analysis of macronutrients shows us concentrations of lipid (F_3,8_ = 1.11, p = 0.4010), protein (*F*_3,8_ = 2.32, p = 0.1520), carbohydrate (*F*_3,8_ = 0.19, p = 0.8950), and glycogen (*F*_3,8_ = 0.51, p = 0.6880) did not differ significantly amongst individuals of the F1, F2, and F3 generations (Table S3 in Supplementary Material).

## 5. Bifurcation diagrams and numerical simulations

In order to parametrize our model, we obtained the survival proportion and fertility parameter estimates from the experimental data by calculating the average number of eggs laid per female individual (*F_i_*) and the average proportion of individuals surviving from one generation to the next (*S_i_*). Our parameter estimates can be seen in Table 1.

For the purpose of simulating model (1), with *g_i_*(*x_i_*) given by (4), we must specify a form for *d_i_*(*x_i_*), as otherwise this is an arbitrary function taking values in (0,1). We will therefore let density-dependent dispersal from patch *i* be given by

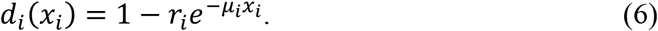

Here *μ_i_* is the strength of density dependence on dispersal and *r_i_* ∈ (0,1) is a scaling factor that can be interpreted as the minimum amount of dispersal that is allowed. If we let the number of individuals that remain on patch *i* follow a Poisson distribution with mean *μ_i_* then *e*^−*μ_i_*^ is the probability of obtaining a zero, i.e., the probability of no dispersal. Hence 1 – *e*^−*μ_i_x_i_*^ may be interpreted as the probability of dispersing from patch *i*, which decreases faster with a growing patch density. An estimate for *α* was obtained from Prout and McChesney (1985), who used a density dependent term of 0.0064 in their demographic model (the sum of f and s, the sensitivity parameters for fertility and survival, respectively). We also set *r*_1_ = *r*_2_ = 0.9, which corresponds to permitting a minimum of 10% dispersal between both patches.

For the dispersal function given by (5) and local dynamics given by (4), we simulated our model for 75 generations, for various combination of μ*_i_* and using the experimentally derived demographic parameters. We assumed a sex ratio of a half, as this accurately reflects the ratio in the models of Prout and McChesney (1985). We can see that after initial increases in both patch densities, trajectories either settle on a stable equilibrium or oscillate around this equilibrium indefinitely. We extended the total number of generations to 500 and these oscillations did not cease but increased in magnitude. Even though both patch populations exhibited oscillatory dynamics, they seem to persist in the long run. Although trajectories approach 0 and then increase rapidly to high densities, these boom-bust type dynamics are common among insect population models (Goodsman, Cook & Lewis, 2017; Strayer et al., 2017).

Different methods have been proposed to analyze the parameter sensitivity in population growth models. Among the approaches used for sensitivity analysis, bifurcation analysis stands out. Bifurcation analysis aims to evaluate the stability properties of equilibria under parameter variations. This is particularly useful for investigating the association of ecological patterns of population oscillations with changing values of demographic parameters, allowing one to quantify this contribution to changes in model outputs (Van Voorn and Kooi 2017).

We conducted a bifurcation analysis to test our model’s stability in the *R_i_F_i_S_i_* and *μ_i_* parameter spaces (see Figure 6). Bifurcation diagrams emerge from relationships between parameter values and population sizes (Dercole & Rinaldi 2012). Usually, the parametric space on one axis determines significant changes on the other axis, expressing long-term population behaviour. As a result of this relationship, it is possible to observe stable trajectories, limit cycles with fixed maximum and minimum limiting values and chaos, a regime characterized by total unpredictability, that is, by the absolute absence of fixed cycles (Luque et al. 2011).

**Figure 6.**
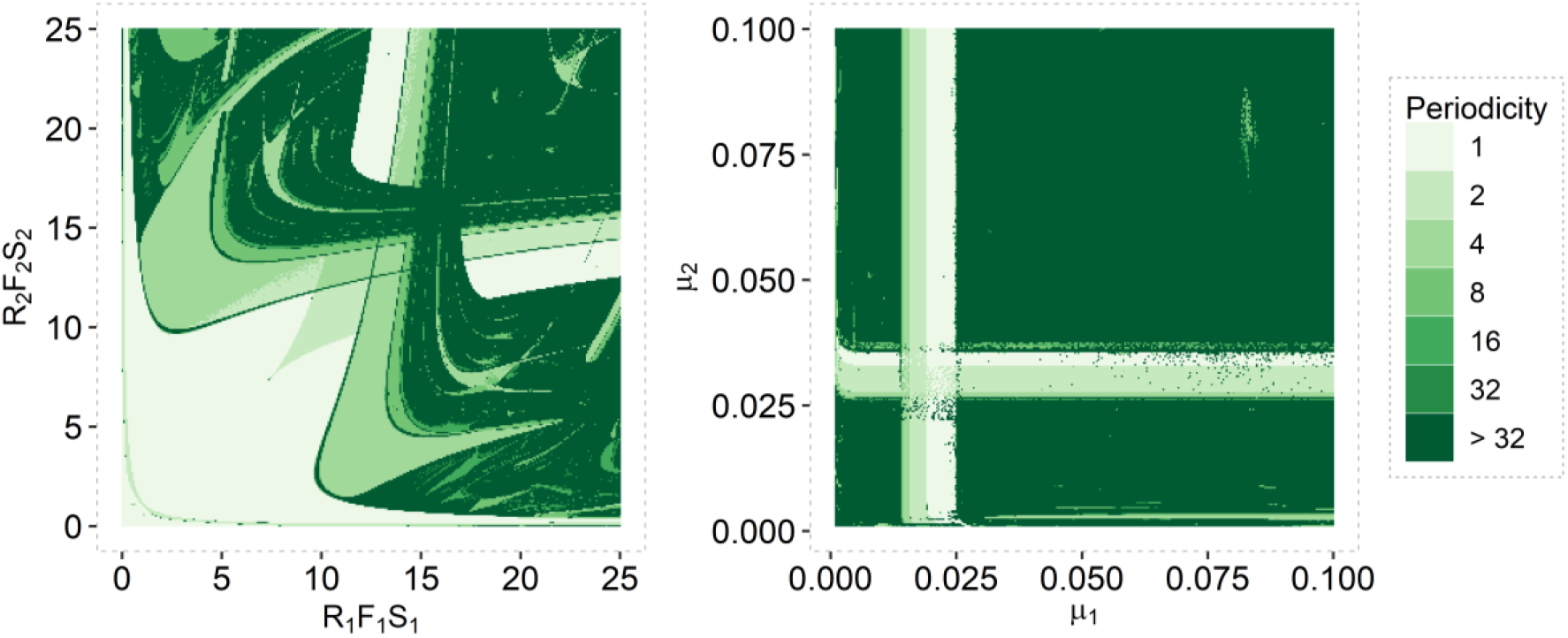
Bifurcation diagrams for the parameter product *R_i_S_i_F_i_* (left) and μ*_i_* (right) for *i* = 1,2. These show the number of unique population sizes according to the colour gradient (stable equilibrium for periods of 1 and periodic orbits for higher values than 1). We considered the last 100 observations after 1000 generations. Initial conditions where (10, 10) for both patches. Note that we only show the bifurcation diagrams for patch 1 as the diagrams are identical for patch 2.

We chose to use the product of the demographic parameters in our bifurcation analysis as they play an important role in determining conditions for persistence and the presence of a positive equilibrium. For the *R_i_F_i_S_i_* bifurcation diagram (Figure 6) we set *μ*_1_ = 0.2 and *μ*_2_ = 0.3, to reflect low to moderate density dependence. We then simulated trajectories for 4 different points that correspond to varying limit cycle periodicities (see Figure 7). In Figure 6 we see that at low values of *R_i_S_i_F_i_* we have stabile equilibria. As we allow demographic parameters to reach values roughly above 10, trajectories are quite unpredictable, in that the period of these limit cycles begin to become greater than 32. Many discrete models exhibit similar complex dynamic behaviour like that seen in Figure 6 and 7 (Hassell, Comins & May, 1991; Geritz and Kisdi, 2004). Trajectories may seem predictable for some parameter ranges, either stabilising or fluctuating after initial increases, but outside these ranges deterministic chaos is generally observed. Note that empirical parameter estimates give *R_i_F_i_S_i_* within the stable region of the parameter space.

**Figure 7.**
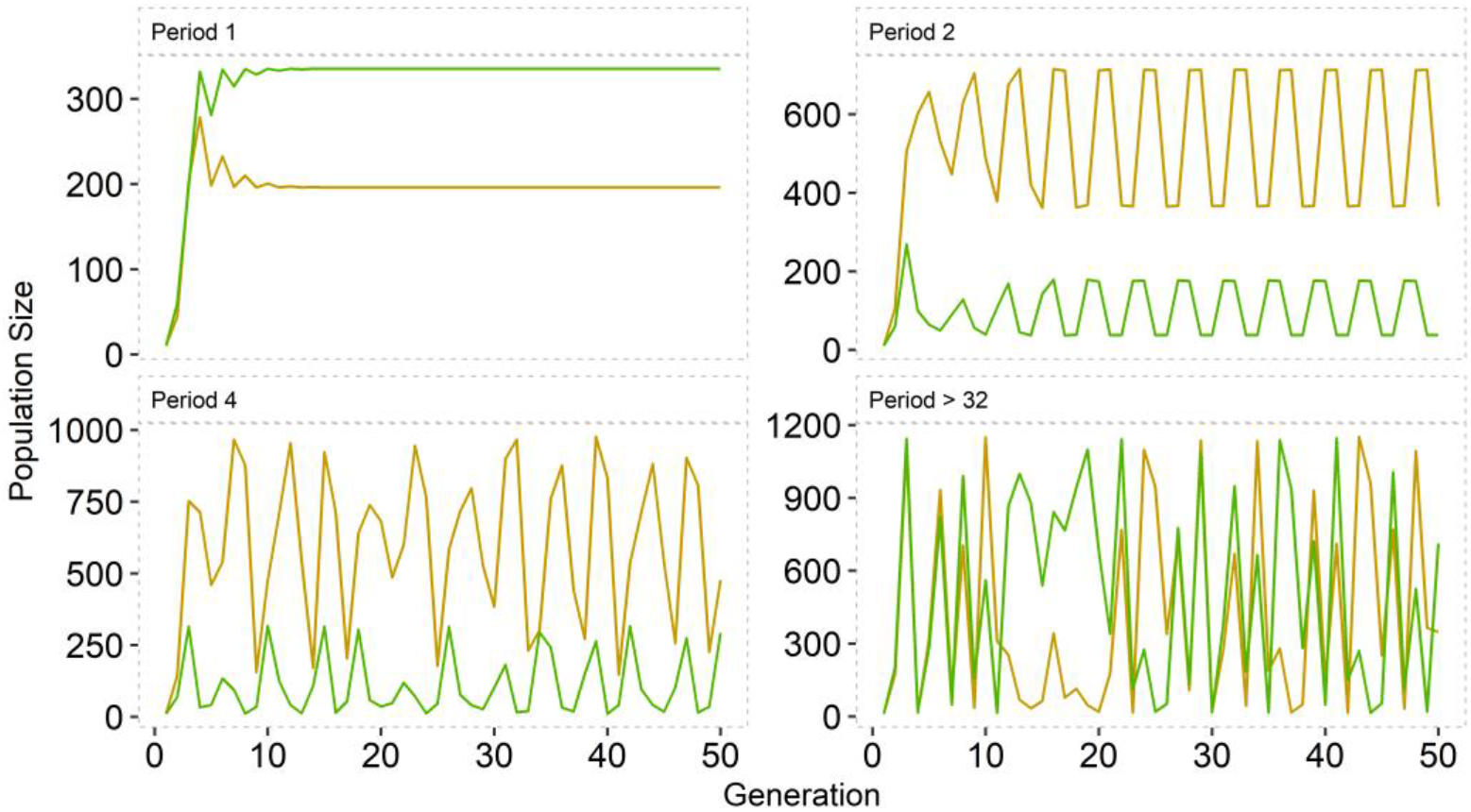
Patch dynamics for patch 1 (yellow) and patch 2 (green) with initial conditions (10, 10), showing the influence of the *R_i_S_i_F_i_* parameter space with μ_1_ = 0.2 and μ_2_ = 0.3. Periods refer to the period of the limit cycles.

For the μ*_i_* bifurcation diagram we set *R*_1_*F*_1_*S*_1_ = 20 and *R*_2_*F*_2_*S*_2_ = 24, as the products obtained from empirical estimates showed stable trajectories for all values of *μ_i_*. Thus, we were interested to explore what happens when both patches exhibit high survival and fertility. The narrow range within the (μ_1_, μ_2_) parameter space (see Figure 6), where trajectories seem to approach a stable fixed point (the uniqueness of which is undetermined), show how sensitive density-dependent dispersal can be. As we increase both parameter values, more complex dynamical behaviour occurs, which may allude to increased density dependent effects on individuals who remain on each patch. This may be because *R_i_F_i_S_i_* is sufficiently greater than 0, meaning subpopulations have higher survival proportions or increased fertility on their respective patches, which can lead to increasing competitive interactions. Each patch also has a nonzero influx/outflow of individuals to/from it, and this may permit sufficient genetic mixing, with dispersal being one of the main drivers of genetic variation in insect species (Raymond et al. 2013).

The demographic parameters considered in Table 2 determine the conditions for the existence of a patch coexistence equilibrium. For values of *μ_i_* within the narrow band in Figure 6 one can observe that we always have *ρ*(***A***(**0**)) > 1. Proposition 2 then implies that we have existence of a positive equilibrium and by Theorems 1 and 2 we have persistence. We have not mathematically explored the stability and uniqueness of this fixed point but instead have derived conditions for our nonlinear population projection matrix that ensures we have both uniform strong persistence and patch-persistence.

## 6. Discussion

The dispersal model employed in this study arose from a classical metapopulation perspective, proposed to analyse the dispersal of insects between finitely many patches (Hanski & Gaggiotti 2004, Moretti et al. 2013, Dey & Joshi 2018). Mathematical models of population growth coupled by migration or dispersal have often been used to analyse population stability by employing the use of bifurcation diagrams (Ruxton 1996; Moretti et al. 2013; Pal et al. 2018). Values of demographic parameters and dispersal rates generally have a strong association with the stability of population equilibria (Moretti et al. 2013; Pal et al. 2018; Cheng et al. 2019).

Dispersal has been suggested as a possible stabilising mechanism for populations that inhabit local sinks, through what has been termed dispersal driven growth (DDG) (Guiver et al. 2017). Our model results further support this hypothesis for spatially structured populations, by showing that dispersal may rescue a species from extinction, even when fertility and survival proportions are low on one of the patches, i.e., one of subpopulations have low local fitness. In model simulations we found that, including density dependent dispersal and intraspecific competition, trajectories exhibit a range of behaviour and are highly sensitive to changes in parameter values. It must be kept in mind that these simulations excluded demographic or environmental stochasticity and so although our final deterministic model ‘skeleton’ seems to approach a positive fixed point after some time, these somewhat smooth dynamics might oscillate significantly, and may even get arbitrarily close to the basin of attraction of the extinction equilibrium (Hastings et al. 2021).

In the context of pest control the opposite of DDG is usually desired (Mazzi and Dorn 2012), and so our results show that the disruption of common dispersal events could be a mechanism to prevent long term population persistence. DDG is an important means for preventing extinction, especially for populations that are dispersing in the absence of external immigration, from matrix habitat or marginal source patches, like for example in agricultural landscapes (Duelli and Obrist 2003). Ephemeral patch dynamics, whereby patches are sequentially restored and destroyed, play a significant role in allowing pest populations that undergo pulsed dispersal to persist (Reigada et al. 2015). Even in the presence of control strategies, such as the use of pesticides or the introduction of natural enemies, pests may exhibit what is referred to as the hydra effect, whereby increasing their mortality rate results in a positive contribution to population growth (McIntire and Juliano 2018; Costa and dos Anjos 2018). We have not considered habitat destruction or inter-species interactions. Extending our work to incorporate these scenarios may prove important, along with increasing the number of patches above the simplest case of two.

Knowing the potential consequences that host shift and biocontrol have on the persistence of crop pests is then vital for food security and for understanding mechanisms of controlling invasions (Bernal and Medina 2018). By including spatial structure in models of herbivore pest dynamics, managers may have the opportunity to determine what series of patches contribute to overall decline/growth of a population (James et al. 2015). Not only does this afford one to know the direct effects of dispersal on a focal host plant, but it also could help one understand the indirect role that pest control could have on adjacent natural matrix habitats (Rand, Tylianakis and Tscharntke 2006). Our models demonstrate that in the absence of dispersal, and under certain parameter constraints, subpopulations on both patches may go extinct. But when connectivity between patches is allowed, we have shown that both uniform strong persistence and patch-persistence can arise, given that the spectral radius of ***A***(**0**) is greater than unity. When this occurs, we have also found the existence of a positive steady state. The stability properties of this fixed point are not known but simulations suggest that populations can either asymptotically approach this (seemingly unique) fixed point, following transient oscillations around it, or oscillate around it indefinitely. For brevity we have excluded an analysis to determine equilibria stability, as persistence is known as a more robust condition that is commonly used to understand a model’s qualitative properties (Jansen and Sigmund, 1998). Ultimately our models suggest that density-dependent dispersal is a vital mechanism by which populations can persevere in a spatial context and inhibiting dispersal, even between two patches (the simplest spatial case), could have significant consequences for the longevity of populations in a patchy landscape.

The bifurcation analysis performed in this study showed regions of stability and instability in the parametric spaces of both the demographic and dispersal parameters. The limit of the values investigated in the bifurcation diagrams are compatible with real values obtained in our experiments. The instability of both populations of *D. suzukii* in general can tend to occur with higher values of the parameters, with exceptions. Narrow ranges that allow for stable trajectories show the importance of parametrising models with experimental data. Among the factors capable of changing the population stability in drosophilids, density, competition, food quality, fecundity and migration stand out (Tung et al. 2018), as demonstrated using our theoretical and empirical findings.

Studies that emphasise the relationship of demographic parameters with the stability of equilibria has long been discussed (Ruxton 1996; Moretti et al. 2013; Pal et al. 2018; Cheng et al. 2019). There are compensatory mechanisms that can explain the relationship between life history and the demographic parameters involved in the growth rate and carrying capacity, which suggests possible compensations for populations in low and high densities, as already investigated in *D. melanogaster* (Mueller & Ayala 1981).

The availability of food resources can determine the formation of high densities per unit of food substrate in insects and consequently trigger intraspecific competition for food (Hardin et al. 2015), which negatively influences the magnitude of demographic parameters associated with population growth and equilibrium (Klomp 1964; Johst; Berryman & Lima 2008). High survival and fecundity values are generally associated with limit cycles and can even produce deterministic chaos (Johst, Berryman & Lima 2008). Polyphagous insects such as *D. suzukii* can resort to fruit exchange because of the fly density on the fruit that has been initially exploited (Hardin et al. 2015; Olazcuaga et al. 2019) and the changing conditions of a food source can thus enhance competition between flies on a per-patch level.

As revised in Dey & Joshi (2018), migration associated with food source can also interfere with the population stability. They studied different numbers of adults migrating between patches and concluded that the effect on persistence and constancy of the population can be altered by food quality and the immigration rate between patches. Considered in an applied scenario, the movement of pests between different patches can result in damaging economic losses. In pest control programs the aim is to present actions that maintain the density of pests in an equilibrium level below damaging economic thresholds (Zhang et al 2018; Wieser et al. 2019; Bryant et al. 2020), so the analysis of parameter values that lead to the loss of stability, resulting in unpredictable population outbreaks, has an unquestionable utility for integrated pest management strategies (Costa & Faria 2010).

In our laboratory experiments the results of the no-choice change of fruits showed that the quality of the food source impacted fecundity, period of oviposition and proportion of viable populations for *D. suzukii*. When exploiting resources in crop edges with different host plants, insects in extensive monocultures have little resource choice. In these circumstances, knowledge of the performance and adaptive potential of invading species is desirable, because successful invaders can become major pests at high densities.

The response of the biological parameters to this host shift showed that disturbances mainly affect the proportion of nutrients necessary to allow immature individuals to develop, as already shown for other *Drosophila* species (Lee, K.P. et al. 2008; Matavelli et al., 2015; Rodrigues et al. 2015). The change in this nutrient proportion, even evaluated indirectly as in the present study, helps to understand the changes in ecological viability of a species. The results for the joint modelling for *D. suzukii*, considering the association parameter, indicate that fecundity can be related to the risk of mortality. Interest in joint models has recently increased, especially for studying the association between longitudinal and time-to-event processes (Alsefri et al. 2020).

In the present study, the high number of eggs of *D. suzukii* found in the second generation, for insects reared on strawberry, may reflect their parental nutritional conditions, as insects of one generation that feed on good-quality host plants are likely to successfully produce subsequent filial generations (Matzkin et al. 2013). The negative impact of biological parameters could not be explained by the intake of macronutrients. A possible explanation is the diversity of the microbiota associated with these insects, since symbiotic bacteria can help their insect host to assimilate nutrients (Shin et al. 2011; Sommer & Newell 2018; Su et al. 2013). The diversity of symbionts can drastically affect nutrient assimilation, which changes with environmental conditions and with corresponding changes to the host food source (Obadia et al. 2018; Yun et al. 2014; Guidolin & Cônsoli 2017), even at optimal nutrient ingestion. Therefore, the development of polyphagous insects may be affected by synergistic factors, including endosymbiosis, as shown for *D. melanogaster* (Shin et al. 2011) and *D. suzukii* (Bing et al. 2018).

## 7. Conclusion

The theoretical and empirical results obtained in this study furthers the understanding of plant host-shift, its possible ecological mechanism, and biological consequences for insect pests. One possible idea that we did not explore but is important to consider, is to include stage structure, such as in the well-known LPA model (Dennis et al. 2001), to deduce what developmental stages may affect plant host-shift and population growth. In order to understand the dispersal dynamics of insect populations, important factors to consider are spatiality and resource availability. Recognizing the explicit movement and within population interactions of insect species will prove to be important for both natural and invading species who inhabit patchy landscapes.

## Data accessibility statement

The reproducible R code and datasets used throughout this paper are available at https://zenodo.org/record/6453797

## Acknowledgements

We thank Dr Daniel Bernardi for kindly providing the first individuals of *D. suzukii* to establish our populations. We thank Ms Felipe Andreazza and Dr Eugênio Eduardo de Oliveira for valuable discussions on the subject, Dr Carolina Reigada for her assistance with the experimental design, and Dr Fernando Luis Cônsoli for help in the macronutrient analyses. The authors thank Dr Janet W. Reid for revising the English text and also thank the Conselho Nacional de Desenvolvimento Científico e Tecnológico (CNPq) for granting a masters scholarship to IBS.

## Supplementary Material

## Appendix 1 - Definitions and Notation

For 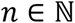 with *n* ≥ 2 denote by 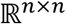 the space of all real positive square matrices of dimension n, and by 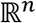 n-dimensional real Euclidean space.

Given an *n* × *n* matrix 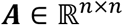, the **spectrum** of **A**, denoted *σ*(***A***), is the set of all eigenvalues of ***A***. The **spectral radius** of ***A*** is defined as

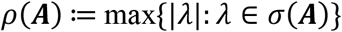

Given a function *F*(***x***) = ***A***(***x***)***x***, for 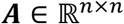 and ***x*** ≥ 0, we denote the Jacobian matrix of *F*(***x***) evaluated at ***a*** as 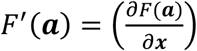.

We write ***x*** ≫ 0 to denote the vector 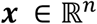 that has all positive entries, i.e. *x_i_* > 0 for all *i* = 1, ⋯, *n*.

For 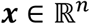 and *p* ≥ 1 we define the *l_p_* vector norm as

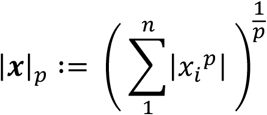

and the matrix norm induced by the *l_p_* norm as

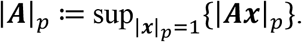

We mainly work with the *l*_1_ norm in this study. We use the same notation |·| for both the vector and matrix norms. This should not cause confusion as we use lowercase letters for vectors and uppercase letters for matrices.

A general nonlinear matrix model with state space 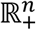 is given by

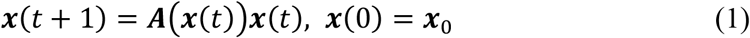

where ***x*** = (*x*_1_ *x*_2_ ⋯ *x_n_*) is a state vector in 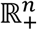 and 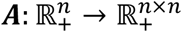 is a nonnegative population projection matrix. The population projection matrix we use in our study is given by

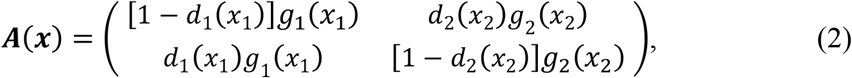

where 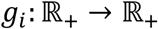 and 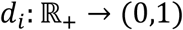 are two continuously differentiable functions that respectively describe local growth on patch *i* and density-dependent dispersal from patch *i*.

An equilibrium ***x***^⋆^ of (1) is a solution of ***x*** = ***A***(***x***)***x***. The equilibrium ***x***^⋆^ is said to be **stable** if for any *ϵ* > 0 there exists a *δ* > 0 such that |***x***_0_ – ***x***^⋆^| < *δ* implies that |***x***(t) – ***x***^⋆^| < *ϵ* for all t, where ***x***_0_ denotes the initial condition ***x***(0). If, in addition, there is some *R* > 0, such that ***x***(*t*) → ***x***^⋆^ as *t* → ∞ for any solution with |***x***_0_| < *R*, the equilibrium is **locally asymptotically stable**. If this holds for any *R* > 0, it is said to be **globally asymptotically stable**.

Given a general nonlinear matrix model of the form (1) with state space 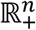 we say that the system is **uniformly weakly persistent** with respect to a persistence function 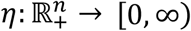 if there exists some *ϵ* > 0, such that for each solution ***x***(*t*) = (*x*_1_(*t*) *x*_2_(*t*) ⋯ *x_n_*(*t*)), with initial condition 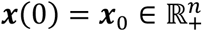 and *η*(***x***_0_) > 0,

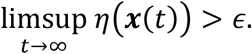

When we replace the *limsup* with the liminf in the above definition we refer to system as being **uniformly (strongly) *η*-persistent**.

We will assume that the local growth function takes the form

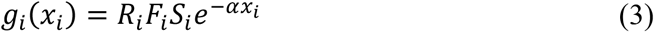

where *F_i_* > 0, *S_i_* ∈ (0,1], *R_i_* ∈ (0,1) and *α* > 0, for *i* = 1,2. Parameters *F_i_*,*S_i_* and *R_i_* can be interpreted as respectively being the fecundity, survival proportion and sex ratio on patch *i*, with *α* being the influence of intra-specific competition for resources on both patches.

A system of the form (1) is said to be **point-dissipative or ultimately bounded** if there is some *M* > 0 such that for all 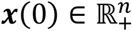, there exists some *T* > 0 such that |***x***(*t*)| ≤ *M* for all *t* ≥ *T*.

Denote by *F^m^* = *F* ∘ *F* ∘ ⋯ ∘ *F* (m times) the m-fold composition of a function 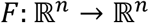 with itself, for *m* a positive integer. A function 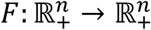 is said to be **strongly positive** if for all *c* > 0 there exists some positive integer *t* > 0 such that *F^t^*(***x***) ≫ 0 for all 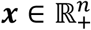 with 0 < |***x***(*t*)| ≤ *c* (Smith and Thieme, 2011).

A matrix 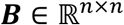 is said to be **irreducible** if there is no proper subset *J* ⊂ {1, ⋯ *n*} for which *b_ij_* = 0 for all *i* ∈ *J, j* ∈ {1, ⋯, *n*} \ *J*.

### Perron-Frobenius Theorem (Horn and Johnson, 1989)

If a nonnegative matrix ***A*** is irreducible, then *ρ*(***A***) > 0 and there exists unique positive right and left eigenvectors corresponding to *ρ*(***A***).

It can be sseen from the above theorem that for 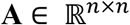 nonnegative, irreducible and n ≥ 2, 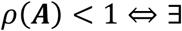 some vector **v** ≫ 0 in 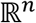 such that ***Av*** ≪ ***v***.

#### Relevant Results from Smith and Thieme (2011)

We recall here the most relevant facts from Smith and Thieme (2011) in this section. Throughout, unless stated otherwise, *F*(***x***) = ***A***(***x***)***x*** is a nonlinear matrix function such as appears in the system definition (1).

##### Theorem A1 (Theorem 7.5 in (Smith and Thieme (2011))

Let 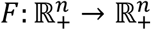 be continuous and assume that there exists *R* > 0, 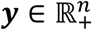 and some nonnegative matrix **D** such that *ρ*(***D***) < 1 and *F*(***x***) ≤ ***y*** + ***Dx***, for 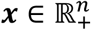 and |***x***| ≥ *R*. Let *F* be differentiable at **0** and further assume that either *F*′(**0**) is irreducible, or *F*(***x***) = ***A***(***x***)***x*** with ***A***(***x***) nonnegative and *ρ*(*F*′(**0**)) > 1. Then there exists some nonzero 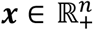 such that *F*(***x***) = ***x***.

##### Theorem A2 (Theorem 7.9 in (Smith and Thieme (2011))

Let 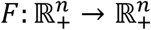 be a differentiable mapping. Suppose that:

1. 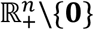 is forward invariant under F.
2. There exists *r*_0_ > 1 and ***v*** ≫ 0 such that 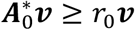, where ***A***_0_ = *F*′(0) and 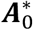 denotes its transpose.
3. The system (1) defined by F is point-dissipative.

Then system (1) is uniformly *η*-persistent for *η*(***x***) = |***x***| where |·| is any norm on 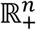.

In the following result, ***x***(*t, **x***_0_) denotes the solution of (1) at time *t* corresponding to the initial condition ***x***_0_.

##### Theorem A3 (Theorem 7.15 in (Smith and Thieme (2011))

Suppose 1-3 from Theorem A2 hold, and F is strongly positive. Let 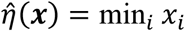. Define

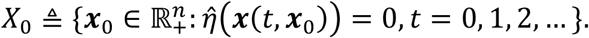

Then there exists some *ϵ* > 0 such that

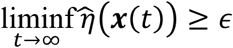

for any solution of (1) with 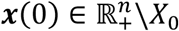.

## Appendix 2 - Proofs

### Proof of Proposition 1

First note that for ***x*** ≥ 0

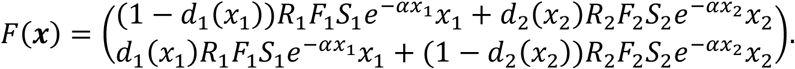

As *x_i_e*^−*αx_i_*^ ≤ (*αe*)^−1^ for *x_i_* ≥ 0, *i* = 1,2, it is easy to see that there exists ***y*** ≫ 0 with *F*(***x***) ≤ ***y*** for all 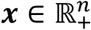. As *F* is continuous and *F*(***x***) = ***A***(***x***)***x***, it follows from Theorem A1 that there exists some 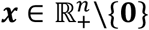 with *F*(***x***) = ***x***. As *F*(***x***) is strictly positive, this implies that ***x*** ≫ 0.

### Proof of Lemma 1

First note that for 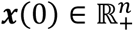 this implies 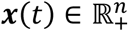 for all *t* ≥ 0. Thus the *l*_1_ norm of ***x***(*t*) is given by **1**^*T*^***x***(*t*). Moreover, for any *t* ≥ 1, ***x***(*t*) = ***A***(***x***(*t* – 1))***x***(*t* – 1) for some 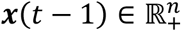. From the form of ***A***(***x***), we can see that for ***x*** = (*x*_1_ *x*_2_)^*T*^,

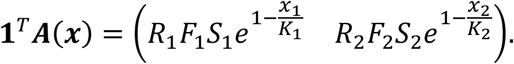

Hence

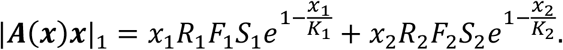

As 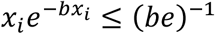, if we let 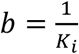 it is clear to see that 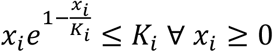.

Therefore

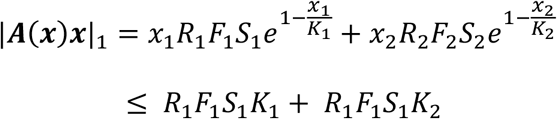

As our solution ***x***(*t*) is of the form ***A***(***x***)***x*** for any *t* ≥ 1, let *M* = *R*_1_*F*_1_*S*_1_*K*_1_ + *R*_2_*F*_2_*S*_2_*K*_2_ and this proves our lemma.

### Proof of Theorem 1

It follows from the form of ***A***(***x***) that ***x*** > 0 implies ***A***(***x***)***x*** ≫ 0. So, it is certainly true that 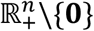 is forward invariant. As *d*(*x_i_*) ∈ (0,1), it can be seen that ***A***(**0**) is irreducible for *α* > 0, *F_i_* > 0, *S_i_* ∈ (0,1] and *R_i_* ∈ (0,1), *i* = 1,2. Also *ρ*(***A***(**0**)) > 1 implies that *ρ*(***A***(**0**)^*T*^) > 1, so using the Perron-Frobenius Theorem there exists some ***v*** ≫ 0 with ***A***(**0**)^*T*^***v*** ≫ ***v***. This implies that we can choose *r*_0_ > 1 such that ***A***(**0**)^*T*^***v*** ≥ *r*_0_***v***. Finally, Lemma 1 implies that the system defined by *F*(***x***) = ***A***(***x***)***x*** is point dissipative. The result now follows immediately from Theorem A2.

### Proof of Theorem 2

*F*(***x***) = ***A***(***x***)***x*** is strongly positive and 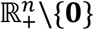 is invariant under *F*. It follows from Lemma 1 that the system is point-dissipative. Note that *F*′(**0**) = ***A***(**0**) is irreducible for *d*(*x_i_*) ∈ (0,1) and for *α* > 0, *F_i_* > 0, *S_i_* ∈ (0,1] and *R_i_* ∈ (0,1), *i* = 1,2. If *ρ*(***A***(**0**)) > 1, then *ρ*(***A***(**0**)^*T*^) > 1 and so there exists some ***v*** ≫ 0, *r*_0_ ≥ 1 such that ***A***(**0**)^*T*^***v*** ≥ *r*_0_***v***, as in the proof of Theorem 1. The result now follows immediately from Theorem A3.

## Appendix 3 - Tables

**Table S1.** Mean oviposition period ± standard error (s.e.) in days, for each generation and host. Means followed by the same letter do not differ statistically.

| Generation | Mean ± s.e. | Host | Mean ± s.e. |
| --- | --- | --- | --- |
| *F1* | 15.20 ± 2.86 a | *Raspberry* | 27.56 ± 2.69 a |
| *F2* | 17.90 ± 3.94 a | *Strawberry* | 5.17 ± 1.20 b |

**Table S2.** Means ± standard errors (s.e.) of the percentage of viable eggs for generations and hosts. Means followed by the same letter do not differ statistically.

| Generation | Mean ± s.e. | Host | Mean ± s.e. |
| --- | --- | --- | --- |
| *F1* | 16.91 ± 5.14 b | *Raspberry* | 21.60 ± 3.49 b |
| *F2* | 32.10 ± 5.88 a | *Strawberry* | 33.47 ± 14.69 a |

**Table S3.** Mean macronutrient concentrations (μg/μL) ± standard errors for each generation.

| Generation | Macronutrient |  |  |  |
| --- | --- | --- | --- | --- |
|  | Lipid | Protein | Carbohydrate | Glycogen |
| *F1 – raspberry* | 10.43 ± 0.1075 a | 0.13 ± 0.0064 a | 0.02 ± 0.0066 a | 0.03 ± 0.0148 a |
| *F2 – raspberry* | 10.37 ± 0.0756 a | 0.13 ± 0.0061 a | 0.02 ± 0.0003 a | 0.04 ± 0.0074 a |
| *F3 – raspberry* | 10.27 ± 0.0563 a | 0.12 ± 0.0049 a | 0.02 ± 0.0014 a | 0.05 ± 0.0123 a |
| *F3 – strawberry* | 10.43 ± 0.0399 a | 0.14 ± 0.0074 a | 0.02 ± 0.0020 a | 0.03 ± 0.0117 a |

